# Recent evolution in *Rattus norvegicus* is shaped by declining effective population size

**DOI:** 10.1101/015818

**Authors:** Eva E. Deinum, Daniel L. Halligan, Rob W. Ness, Yao-Hua Zhang, Lin Cong, Jian-Xu Zhang, Peter D. Keightley

**Affiliations:** Institute of Evolutionary Biology, University of Edinburgh, Edinburgh, United Kingdom; State Key Laboratory of Integrated Management of Pest Insects and Rodents in Agriculture, Institute of Zoology, Chinese Academy of Sciences, 1# Bei-Chen West Road, Beijing 100101, China; Institute of Plant Protection, Heilongjiang Academy of Agricultural Sciences, Harbin 150086, China

**Keywords:** *Rattus norvegicus*, evolutionary adaptation, comparative population genomics, effective population size, bottleneck, distribution of fitness effects, PSMC, DFE-α, ILLUMINA whole genome sequencing, *Mus musculus castaneus*

## Abstract

The brown rat, *Rattus norvegicus*, is both a notorious pest and a frequently used model in biomedical research. By analysing genome sequences of 12 wild-caught brown rats from their ancestral range in NE China, along with the sequence of a black rat, *R. rattus*, we investigate the selective and demographic forces shaping variation in the genome. We estimate that the recent effective population size (*N_e_*) of this species = 1.24 × 10^5^, based on silent site diversity. We compare patterns of diversity in these genomes with patterns in multiple genome sequences of the house mouse (*Mus musculus castaneus*), which has a much larger *N_e_*. This reveals an important role for variation in the strength of genetic drift in mammalian genome evolution. By a Pairwise Sequentially Markovian Coalescent (PSMC) analysis of demographic history, we infer that there has been a recent population size bottleneck in wild rats, which we date to approximately 20,000 years ago. Consistent with this, wild rat populations have experienced an increased flux of mildly deleterious mutations, which segregate at higher frequencies in protein-coding genes and conserved noncoding elements (CNEs). This leads to negative estimates of the rate of adaptive evolution (*α*) in proteins and CNEs, a result which we discuss in relation to the strongly positive estimates observed in wild house mice. As a consequence of the population bottleneck, wild rats also show a markedly slower decay of linkage disequilibrium with physical distance than wild house mice.

## Introduction

Comparative genomics became possible between human and mouse with the publication of the mouse genome (Mouse Genome Sequencing Consortium, 2002), leading to many important new findings, including estimates of the fraction of conserved nucleotide sites, corroboration of downwardly revised estimates of protein-coding gene number, and the discovery of ultraconserved non-coding elements (Bejerano et al., 2004). Genome sequencing of single individuals naturally led to population genomics, pioneered in *Drosophila simulans* (Begun et al., 2007). This allows detailed inferences to be made concerning many important questions in evolutionary genetics, including the demographic history of populations (Li and Durbin, 2011), the nature and frequency of adaptive evolution (e.g., Hernandez et al., 2011; Sattath et al., 2011), and the causes of the correlation between recombination rate and neutral diversity (Cai et al., 2009).

Much population genetics has traditionally relied on comparing nucleotide polymorphism in one species to divergence from another. With decreasing sequencing costs, comparative population genomics - the comparison of multiple genome sequences from different species - has now become possible. This allows reciprocal analysis of polymorphism versus divergence. For example, the McDonald-Kreitman test (McDonald and Kreitman, 1991) for estimating the relative extent of adaptive evolution, uses polymorphism in one species (A) and the divergence between two species (A and B). From the assumptions of the test, including a stable demography since the divergence between the two species, the reciprocal estimate should yield the same result. A significant difference in the estimates could, therefore, indicate that an evolutionary important demographic change has occurred since the split. For such a reciprocal analysis to be biologically meaningful, the species need to be closely related, but not so closely related that they share a substantial fraction of nucleotide polymorphism that originated prior to the split between the species. This condition is comfortably met in the case of our focal species, wild brown rats and mice, which are thought to have diverged at least 12MYA (Benton and Donoghue, 2007), equating to at least 24 million generations.

One of the primary determinants of the efficacy of selection across the genome is the strength of genetic drift. The rate of drift is inversely proportional to the effective population size (*N_e_*), which represents the size of an ideal population that would display the observed amount of drift. *N_e_* predicts a number of fundamental properties of natural populations, including the amount of genetic variation (*N_e_*× mutation rate (*µ*)), the rate at which linkage disequilibrium (LD) is broken down (*N_e_*× recombination rate (*r*)) and the strength of selection (*N*_*e*_× selection coefficient (*s*)). For example, theory predicts that if the scaled selection coefficient is below one (*N_e_s* < 1) genetic drift will dominate over selection, rendering such mutations ‘effectively neutral’. The proportion of such mutations is therefore predicted to increase across the genome through demographic processes that reduce *N_e_*, such as fluctuating population size or bottlenecks. The interaction between genetic drift and selection can be manifest in a number of different ways, including an increase in the fraction of mutations in functional regions that behave as slightly deleterious or a lower rate of adaptive evolution.

In this paper, we compare a population genomic dataset of wild brown rats with a previously published dataset from wild house mice (Baines and Harr, 2007; Halligan et al., 2013) and investigate the differential effects of drift on the genomic signature of selection acting in protein-coding and conserved non-coding DNA in the genome. The effective population size in our focal population of wild house mice (*Mus musculus castaneus*), is nearly two orders of magnitude higher than recent *N_e_* for human populations and substantially higher than that of inbred lab strains (Salcedo et al., 2007; Baines and Harr, 2007; Phifer-Rixey et al., 2012). Halligan et al. (2013) inferred that the protein-coding and conserved noncoding elements (CNE) evolved more rapidly than the neutral theory expectation, suggesting that there has been substantial genome-wide adaptation in proteins and CNEs of wild mice. Moreover, they found reductions in neutral diversity around protein-coding exons and CNEs, indicative of frequent selective sweeps and/or background selection. In contrast, despite a presumably large contemporary census population size in wild brown rats (*Rattus norvegicus*), it has been estimated that their *N_e_* is five-fold smaller than wild house mice (Ness et al., 2012). The difference in the *N_e_* between mice and rats provides an opportunity to investigate the effects of genetic drift in the mammalian genome and the way in which selection and drift interact to shape patterns of diversity in the genome.

Using whole genome data from 12 rats collected from their ancestral range in NE China and a comparable dataset in wild house mice, we ask a number of questions (1) Does reduced *N_e_* in rats lead to reduced efficacy of selection on new mutations affecting protein or CNE sequence? (2) How does the effect of hitchhiking differ between mice and rats and how does this compare between protein-coding exons and conserved non-coding elements? (3) Does reduced *N_e_* in rats influence the extent of LD? (4) What can patterns of DNA polymorphism tell us about the recent demographic history of wild brown rats? We find strong evidence for a population bottleneck that has distorted the distribution of allele frequencies throughout the genome and altered patterns of LD in wild rats. We also find evidence for a higher frequencies of segregating deleterious mutations in wild rats, consistent with a reduction in the efficacy of purifying selection. However, neutral diversity reductions around protein-coding exons follow a virtually identical pattern in the two species, suggesting that selection has had similar effects on diversity at sites linked to exons, and that these patterns are insensitive to recent changes in *N_e_*.

## Results

### Inference of selective forces operating in protein-coding genes and CNEs

As a first step to quantify and compare the selective forces acting on variation in the wild rat genome, we computed nucleotide diversity (*π*) within brown rats, nucleotide divergence (*d*) from the house mouse and black rat, and Tajima’s *D* for different classes of sites. We focused on two classes of conserved sequences: protein coding exons and CNEs. In exons, 0-fold degenerate sites are under the strongest selection, as any nucleotide change would result in an amino acid substitution. The 4-fold degenerate sites, on the other hand, are typically seen as a neutral standard for these, as mutations at these sites do not result in amino acid changes. CNEs are not translated to proteins, so we consider all sites within CNEs as potentially under selection for the purpose of our analysis. As a neutral standard we use CNE flanking sequence at a distance of at least 500 bp with the same total length as the element (following Halligan et al., 2013).

Both nucleotide diversity and divergence from the outgroups reflected this expected ranking of selection strength (fig. 1, supplementary table S1, see also figs. S1, S2, S3, S4). Diversity was lowest at 0-fold sites (0.045%), followed by CNEs (0.097%), 2-fold sites (0.112%), 4-fold sites (0.147%) and highest in CNE flanks (0.157%). Divergence from the mouse reference sequence (mm10) followed the same rank order (3.1%, 7.1%, 10.9%, 14.2% and 15.4%). In line with this, Tajima’s *D* was similar for 0-fold sites and CNEs (−0.43 and −0.41, respectively) and values for these sites were lower than at 2-fold sites, 4-fold sites and CNE flanks (−0.27, −0.23 and −0.22, respectively). Although all of these values are negative (implying a slight excess of low-frequency variants), they are much closer to zero than what was previously found in wild mice. A recent population bottleneck would affect the genome wide diversity spectrum in a way that produces less negative, or even positive, Tajima’s *D*. Additionally, *π*/*d* was higher for 0-fold sites and CNEs than for the other classes (0.015 (0-fold), 0.014 (CNEs), 0.010 (other classes)). This pattern is expected if there are slightly deleterious mutations affecting 0-fold sites and CNEs, since they are expected to contribute more to *π* than to *d*.

**Figure 1.**
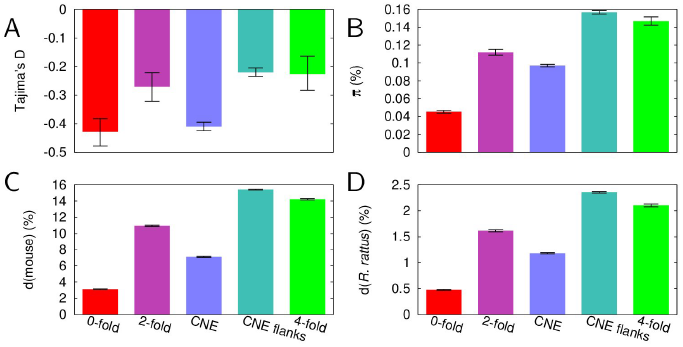
Tajima’s *D* (A), diversity (B) and divergence from mouse (mm10; C) and *R. Rattus* (D) for wild *R. norvegicus* data. Error bars show 95% confidence intervals based on 1000 bootstrapping replicates. See supplementary material for impact of proximity filter, exon/CNE selection rules (figs. S1, S2, S3 and S4) and restricting analysis to sites with corresponding R. rattus bases (table S1)

We then estimated distributions of fitness effects (DFE) of new mutations using the program DFE-*α* (Keightley and Eyre-Walker, 2007) and compared the results to previous estimates from wild house mice (Halligan et al., 2013) (fig. 2). DFE-*α* compares the folded site frequency spectra (SFSs) of two classes of sites, one neutrally evolving and one under selection, to assess the DFE. It uses that mutations are purged faster from the selected sites when they are more deleterious. In line with theoretical expectations for a smaller *N_e_* in the rat, we inferred that there is a substantially larger proportion of mildly deleterious mutations (*N_e_s* < 1), 0.29 and 0.58 in exons and CNEs, respectively, than in the same classes in wild mice (0.17 and 0.44, respectively). Concordantly, the proportions of highly deleterious mutations (*N_e_s* > 10) were lower in the rat (exons: 0.65 and CNEs: 0.29) than in the mouse (0.77 and 0.37, respectively). The presence of a higher fraction of slightly deleterious mutations is consistent with the increased values of *π*/*d* that we found for exons and CNEs.

**Figure 2.**
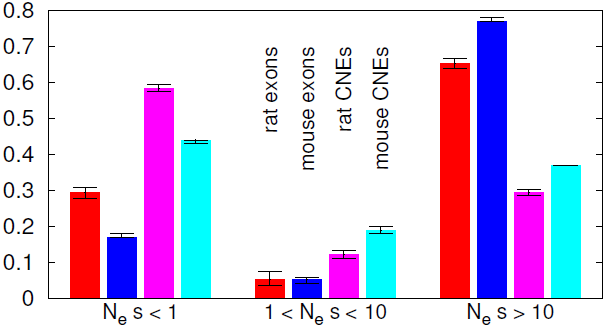
DFE of deleterious mutations for rat exons (red), mouse exons (blue), rat CNEs (magenta) and mouse CNEs (cyan). Error bars indicate 95% confidence intervals from 1000 bootstrapping replicates. Mouse data after Halligan et al. (2013)

We also attempted to estimate the fraction of nucleotide differences in exons and CNEs driven to fixation by positive selection (*α*) and the rate of adaptive substitution relative to the rate of neutral substitution (*ω_α_*). For this we used an extension of the McDonald-Kreitman test incorporated into DFE-*α* (McDonald and Kreitman, 1991; Eyre-Walker and Keightley, 2009). This subtracts the number of fixed nucleotide substitutions (relative to the mouse) in the selected class of sites that is expected from the fixation of neutral and deleterious mutations alone from the actual number of substitutions. The remainder is contributed to positive selection. We consistently obtained negative estimates for *α* and *ω_α_* for both exons (table S2). As we discuss below, these negative estimates likely reflect a recent population size bottleneck.

### Reduced diversity around exons and CNEs

Selection operating within CNEs and exons is also expected to affect nucleotide diversity in closely linked surrounding sequences as a consequence of selective sweeps (Maynard Smith and Haigh, 1974) or background selection (Charlesworth et al., 1993). We therefore investigated diversity statistics in exonic (up to 100 kb) and CNE (up to 20 kb) flanking regions, and again compared our results with those previously obtained in wild house mice (Halligan et al., 2013) (fig. 3). Although wild house mice have much higher diversity than what we observe in the rat, the relative reduction in diversity in exon flanks was remarkably similar in both species. The reduction in diversity (*π* and *π*/*d*) around CNEs, on the other hand, was less pronounced in rat. Moreover, in the bins directly adjacent to the CNEs, we found an increase of *π*/*d*, coinciding with a less strong reduction of *π* than in mouse at these strongly conserved sites.

**Figure 3.**
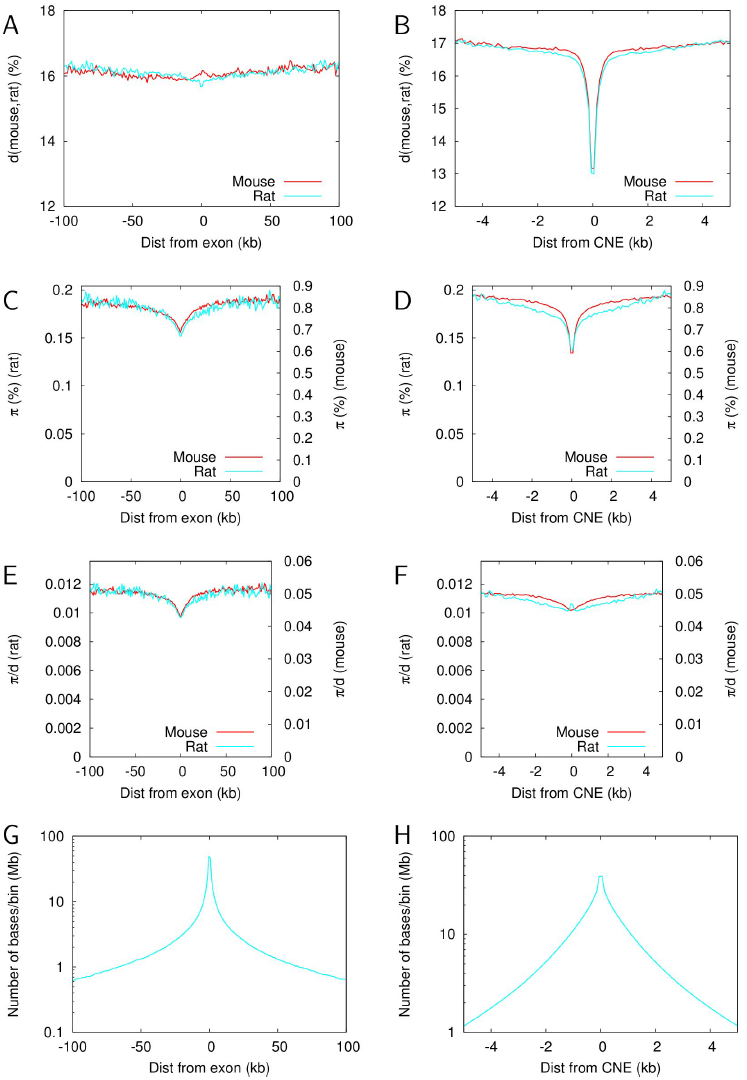
Exon (A, C, E, G) and CNE (B, D, F, H) flanks. A,B: divergence between rat and mouse. C,D: Diversity (π). To facilitate comparison of relative changes between both species, rat values are indicated on the left axis, mouse values on the right axis. E,F: π/*d*. G,H: Number of sites per bin. Each site was only counted once, in relation to its nearest exon/CNE. We used a bin size of 1 kb for exon flanks (up to 100 kb) and 100 bp for CNE flanks (up to 5 kb). Rat data in cyan, mouse data in red. We assessed the impact of the proximity filter (with md=5) on nucleotide diversity in exon and CNE flanks. This resulted in an overall reduction of π of ∼17% and some increase of the π/*d* spike directly surrounding the CNEs, without further changing the general shape of the curves (fig. S16). The magnitude of this reduction was 1-2 times as large as the proximity filter’s impact within exons (0-fold: 17%, 4-fold: 9%) or CNEs (11%).)

The reduction of *π* and *d* in the CNE flanks appears to exist on two length scales: a strong reduction that decays over ∼500 bp and a second less pronounced reduction that decays over tens of kb. We therefore investigated to what extent this second length scale could be the result of the proximity of CNEs to exons. We first computed the distribution of distances from CNEs to the nearest exon (figs. S5, S6). Due to the power law-like distribution of distances between neighbouring exons, bases tend to “cluster” around exons; this means that more bases are located as a particular short (e.g. 10 bp) distance from the nearest exon than at a particular large distance (e.g. 100 kb) from it (fig. 3G). Taking this into account, there remains a two-fold overrepresentation of CNEs near exons (fig. S5). Yet when we used the distribution of distances from CNEs to their nearest exon (fig. S6) to convolute (≈ blur; see methods) the exon flanks, the resulting slope “far away” from the CNEs, i.e., fitted between 5 and 20 kb away, was much shallower than the long length scale in the CNE flanks (fig. S7). This implies that the proximity to exons of many CNEs can only explain a small part of the long length scale we observed in the decay of *π* in CNE flanks.

### LD decay in rat and mouse genomes

To gain an understanding of the striking similarity of the diversity reductions in the exon flanks between rat and mouse and the less similar patterns in CNE flanks, we investigated the decay of linkage disequilibrium (LD) around focal SNPs in wild rats and mice. For this, we computed the pair-wise genomic *r*^2^ (Rogers and Huff, 2009), and averaged over all SNPs at a particular distance from each focal SNP (using a bin size of 20 bp). As focal SNPs, we used either all SNPs, SNPs within exons or SNPs within CNEs (fig. 4AB).

**Figure 4.**
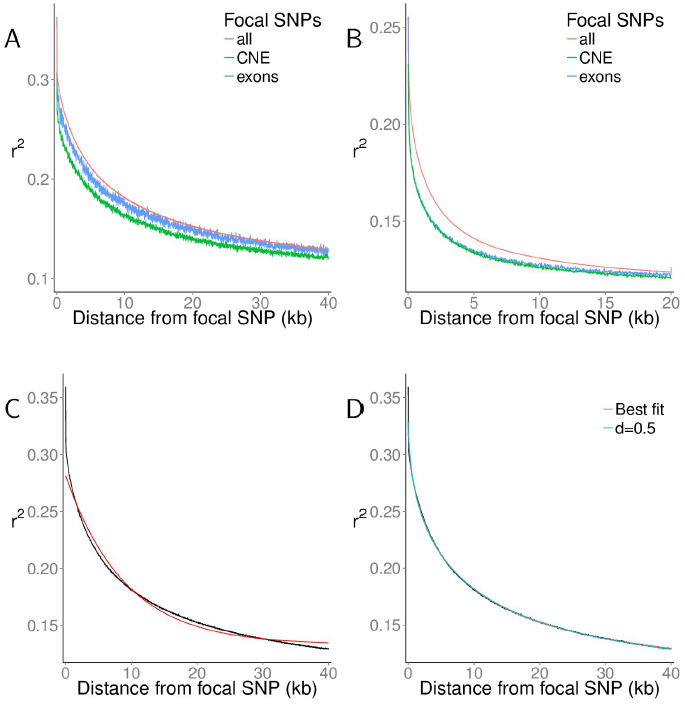
LD measured by genomic 〈*r*^2^〉 as a function of distance from focal SNP. A,B: 〈*r*^2^〉 for all chromosomes combined, with all SNPs, or only those within exons or CNEs as focal SNP. A: rat, B: mouse. All values are averages over 20 bp bins. C: Average of *r*^2^ for all chromosomes combined, all rat SNPs, with offset exponential fit *f*(*x*) = (*a* − *c*) * exp(−*x*/*b*) + *c*. D: Same curve, with an offset stretched exponential fit: *g*(*x*) = (*a − c*)*\** exp(−*(x/b*) ) + c.

In wild house mice, average *r*^2^ (written 〈*r*^2^ 〉) decayed much faster than in wild rats, and the peak value was lower, consistent with the larger *N_e_* in mice. To quantify the difference, we first fitted exponential functions to the decay of 〈*r*^2^〉 with physical distance: *f*(*x*) = (*a* − *c*) × exp(−*x*/*b*) + *c*. For our purposes, characteristic length *b* is the biologically most important parameter, as this is the distance over which the relevant information, 〈*r*^2^〉 − *c*, decays by a factor 1/e (to ∼37% of its original value). Maximum value *α* is the intercept and *c* is the offset, which has a theoretical minimum of 1/(*n*−1) for a sample of *n* individuals (see supplementary text 2) and increased with decreasing *N_e_* (Hill, 1981).

We found that it was impossible, however, to obtain a good fit with this kind of curve (fig. 4C). The structure of the residuals of the best fitting curves (fig. S8AB) suggested that LD (〈*r*^2^〉) decays first faster, then slower than exponential, which is a property of a stretched exponential *g*(*x*) = (*a*−*c*) ×exp(−(*x*/*b*)^*d*^)+*c*, with stretching exponent 0 < *d* < 1. See discussion for the biological meaning of *d*. We obtained good fits to the data with this formula (figs. 4D, S9, table S3A). The rat:mouse ratios of *c* − 1/(*n* − 1) were all larger than 1, consistent with a larger *N_e_* in mouse.

We fitted all curves again with a fixed stretch exponent of *d* = 0.5 to allow a more direct comparison of the characteristic length parameters *b* (figs. 4D, S9, table S3B). This effectively exploits the fact that stretched exponentials are notoriously hard to fit to our kind of data. By these means, we found that LD decays 6-7 times faster in mouse than in rat (rat:mouse ratios of *b*: 7.14 exons; 6.31 CNEs “noOverlap”; 5.96 CNEs “strict”; 5.76 all SNPs).

### Recent bottleneck in the rat population

The preceding analysis provides several lines of evidence for a recent population size bottleneck in wild rats: the much lower diversity than in wild mice (figs. 1B, 3), the negative estimates of *α* and *ω_α_* (table S2), the values of Tajima’s *D* (fig. 1A) that are much larger than in mice and near zero for some functional categories of exons.

To further investigate the possibility of changes in population size, we used the method of Li and Durbin, PSMC, (Li and Durbin, 2011) to infer population history based on the length distribution of genome stretches that are identical by state. Based on the non-CpG prone sites, the 12 rat genomes showed a 3-fold decline in *N_e_* up to 10,000 – 20,000 YA (fig. 5).

**Figure 5.**
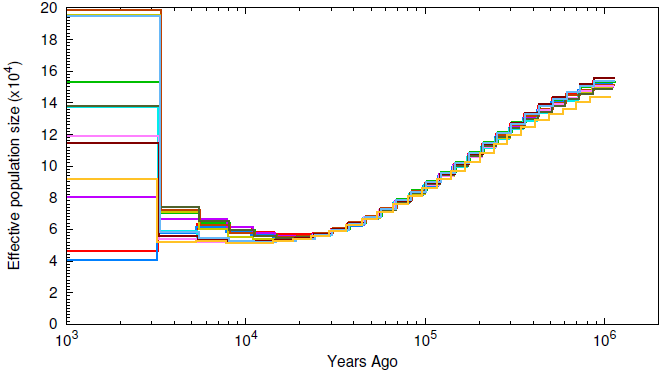
Inferred population history from distribution of IBS (identical by state) tract length distributions Li and Durbin (2011). Curves represent the inferred population history of individual rat samples. Estimates are based on non-CpG prone sites only and using a proximity filter of md = 5 bp. see figs. S10 and S11 for effects of these choices.

A similar trend was found using all sites (i.e., including the CpG-prone sites and adjusting mutation rates accordingly; fig. S10), using a different published mutation rate for rat of *µ* = 4.2 x 10^−9^ (Ness et al., 2012) (fig. S11B) and by varying the parameter (md) of the proximity filter (fig. S11), which is the filter that had most impact information used by PSMC, i.e., the distribution homozygous runs (HHn from MacLeod et al., 2013, see fig. S12). For more details, see supplementary text 3.

From this we conclude that the rat genomes contain a strong and robust signal for a bottleneck between 10,000 and 50,000 YA. The actual bottleneck may have been sharper than the PSMC traces show – and, if it has been fairly short, also more severe – as simulations have shown that PSMC has a tendency to smoothen sharp transitions (Li and Durbin, 2011).

### Discussion

Previous work suggests that wild brown rats have a much lower effective population size than wild house mice (Ness et al., 2012). Re-estimating the mutation rate based on our measured divergence from mouse at 4-fold degenerate sites (14.2%), assuming a divergence time of 12 MYA (Benton and Donoghue, 2007) and 2 generations per year (Halligan et al., 2013; Ness et al., 2012), yields *µ* = 2.96 × 10^−9^(2.94 × 10^−9^ − 2.98 × 10^−9^). Equating *π* at 4-fold degenerate sites to 4*N_e_µ*, we obtain an estimate of *N_e_* = 1.24 × 10^5^(1.20 × 10^5^ − 1.28 × 10^5^), marginally smaller than the 1.3 × 10^5^ estimate previously published. We obtained a very similar *N_e_* estimate using divergence and diversity from our other neutral class, the CNE flanking sites: *N_e_* = 1.22 × 10^5^ (1.21 × 10^5^ − 1.23 × 10^5^) (and *µ* = 3.21 × 10^−9^ (3.20 × 10^−9^ − 3.23 × 10^−9^)).

From our PSMC demographic inference we obtain a minimum *N_e_* of approximately 4 × 10^4^ at 2 × 10^4^ years ago and a 3-4 times larger ancestral size (i.e., the most ancient *N_e_* that can be detected with this method). The *N_e_* estimate based on silent site diversity lies between these values, which is likely explained by the fact that it is affected by the whole recent population history. An effective population size of 4 × 10^4^ appears small for the present day, given the abundance and widespread distribution of brown rats, and the small amount of population structure that appears to be present in wild rat brown populations (Ness et al., 2012). However, it is likely that the effective size of the brown rat population has increased dramatically since the origins of human agriculture, which provided them with a large and stable source of food. None of the methods we have used, however, can reliably estimate contemporary *N_e_*.

### Negative estimates of the rate of adaptive molecular evolution

Conceptually, the rate of adaptive molecular evolution should always be a positive number. We now review the computational method to explain how a recent population bottleneck – as suggested by the unusually high values of Tajima’s *D* (fig. 1) and the PSMC demographic inferences (fig. 5) – could cause negative estimates of *α*, the fraction of substitutions driven to fixation by positive selection. *ω_α_*, the rate of adaptive substitution relative to the rate of neutral substitution, is defined such that it inherits the sign of *α* (Eyre-Walker and Keightley, 2009) OR (Halligan et al., 2013).

To estimate *α* (and *ω_α_*), DFE-α subtracts an estimate of the fraction of (slightly) deleterious substitutions in the selected class (e.g., 0-fold sites) from the total number of substitutions in the selected class, and assumes that the remaining substitutions have been positively selected. To estimate the (slightly) deleterious fraction, the inferred DFE is used along with a single effective population size (weighted over the whole evolutionary time period under analysis), and the selected (e.g., 0-fold) and neutral (e.g., 4-fold) divergence from an outgroup. This implicitly assumes that the strength and effectiveness of selection (i.e., the DFE) were constant over the evolutionary time period under analysis (Eyre-Walker and Keightley, 2009). A recent population size bottleneck is expected to cause an over-representation of slightly deleterious mutations in the polymorphism data used to infer the DFE, which will therefore increase the predicted divergence in the selected site class. This implies that the DFE-*α* method, and all other similar implementations of the McDonald-Kreitman test, will tend to under-estimate the rate of adaptive evolution if there has been a recent population bottleneck. At the heart of this problem lies the fact that divergence is built up continually from the moment the two lineages split (in this case 12 MYA), whereas polymorphism only reflects a limited window of recent evolutionary history (4*N_e_* generations). As the size of this window is dependent on *N_e_*, pre-bottleneck information is rapidly lost from polymorphism data as a consequence of a severe bottleneck. Analysing data from species with shorter divergence times may mitigate the impact of long term population size changes, albeit at the expense of power. Using *R. rattus* as an outgroup could not mitigate the problem, as the split between both rat species is ∼2.9 MYA (Robins et al., 2008), i.e., long before the bottleneck we inferred from our data. Consistently, we again obtained negative estimates of *α* and *ω_α_* (table S2).

### Decay of LD with physical distance

We inferred that there is a roughly 6-7 fold faster LD decay in wild mice than wild rats as a function of physical distance in the genome by fitting stretched exponential functions of the form (*a* − *c*) × exp(−(*x*/*b*)^*d*^) + *c*. This is consistent with a recent population size bottleneck, as confirmed using different information such as our PSMC analysis. We found that a stretched exponential including an offset was needed to obtain a satisfactory fit to the data, whereas a single exponential gave a poor fit (figs. 4, S9).

A stretched exponential can be obtained by summing over a large number of single exponentials with various exponents (parameter *b* in our functions), (e.g., Johnston, 2006). Assuming that the recombination rate is constant over the genome, population genetic theory predicts that LD (〈*r*^2^〉) decay follows a single exponential (Charlesworth and Charlesworth, 2010), which can be offset by the theoretical minimum for the sample size (1/(*n* − 1), see supplementary text 2) + a residual offset for the expected genome wide LD due to finite population size (Laurie-Ahlberg and Weir, 1979; Hill, 1981). The single exponential would also fit if fluctuations in recombination rate average out over sufficiently short distances.

Previous estimates of the recombination rate in rat and mouse (Jensen-Seaman et al., 2004), however, showed evidence of fluctuations on a scale of much more than 10 Mb, i.e., larger than our window for calculating 〈*r*^2^〉, and also showed evidence for variation in whole chromosome average recombination rate. This variation could explain the requirement for a stretched exponential. As the major fluctuations in the reported data for mouse and rat are on the same length scale, this also explains that we found smaller values of the stretch exponent d for mouse than for rat (table S3A): the smaller the relevant window for LD decay, the larger the variation in the (weighted) average recombination rate over the window. The roughly 6-7 fold faster LD decay in mouse also explains why the relative differences among classes (exons, CNEs, all data) of the fitted values are larger in mouse.

In principle, the fitted value of the offset c minus its theoretical minimum carries information about *N_e_* (Laurie-Ahlberg and Weir, 1979; Hill, 1981). With our window length and limitations on resolution close to the focal SNP, however, it is impossible to obtain a reliable estimate of the stretch exponent *d* or the offset c independently. We, therefore, refrain from estimating *N_e_* in this way.

### Reductions of nucleotide diversity around protein-coding exons and CNEs

A striking finding from our study is the extremely similar proportional reductions in mean scaled neutral nucleotide diversity around protein-coding exons in wild rats and in wild house mice (fig. 3). The depth, width and shape of the reductions in diversity are all similar. The drops in diversity are presumably caused by the hitchhiking effect of selection on variants in protein-coding exons, which reduces diversity in tightly linked flanking regions. The previous analysis of the pattern of diversity reduction in wild house mice suggests that there is a substantial role for selection of advantageous mutations, whereas a background selection model alone appears to be incapable of explaining the width of the observed mean diversity reduction (Halligan et al., 2013). The question then arises as to whether these similar patterns around exons in the two species can be reconciled with the difference in the effectiveness of selection between the species (caused by lower *N_e_* in rats) and the presence of substantially greater LD in rats than in mice. If diversity at linked sites is reduced by selection of newly arising advantageous mutations in exons of large selective effect(s) that go to fixation in both species (i.e., classic selective sweeps such that *N_e_s* ≫ 1), and the rate and strength of advantageous mutations and the rate of recombination per physical distance are the same in the two species (0.555 cM/Mb in rat and 0.528 cM/Mb in mouse (Jensen-Seaman et al., 2004)), then equivalent patterns of diversity reduction are predicted, and these are not expected to depend on *N_e_* or LD (Maynard Smith and Haigh, 1974). A similar argument can be made for the case of background selection (BGS) involving strongly deleterious mutations (Nordborg et al., 1996). Alternatively, if diversity reductions are caused by positive selection on standing variation, the pattern of diversity reduction is expected to depend on the effective population size during the phase in which a variant can rise to a high frequency by drift, and subsequently be positively selected to fixation (Przeworski et al., 2005). Specifically, a higher *N_e_* increases the difference between the pattern of diversity change seen under a classic sweep model and a model of standing variation. The similarity of the diversity reduction patterns surrounding protein-coding exons we observe between mice and rats, species which differ substantially in recent *N_e_*, is therefore indirect evidence in favour of the classic selective sweeps model.

Narrower and shallower scaled diversity reductions in the regions surrounding CNEs are also present, but these have somewhat different patterns in mice and rats (fig. 3). Specifically, diversity reductions are shallower in rats and diversity returns to a genomic background level more slowly in rats than mice. It was previously shown in mice that the diversity reductions can be explained by a BGS model, although a role for positive selection was not excluded (Halligan et al., 2013). If diversity reductions are mainly caused by BGS, a weaker effect is expected in rats than mice if there are substantial numbers of CNE mutations which have selective effects < 1/*N_e_* in rats and > 1/*N_e_* in mice, because these would behave as nearly neutral in rats and therefore have a smaller influence on linked neutral diversity than in mice. This is consistent with our estimates of the DFE in the two species, which suggest that there are substantially more deleterious mutations in CNEs with selective effects < 1/*N_e_* in rats than mice (fig. 2).

## Conclusion

We have conducted a whole genome polymorphism study to quantify the selective forces shaping recent wild brown rat evolution and compared our findings to a similar study in wild house mice. We found a larger proportion of slightly deleterious mutations in rats than in mice for both protein-coding exons and CNEs, in line with the theoretical expectation for a larger *N_e_* in mice. The data also shows evidence for a recent population bottleneck in rats, which we dated at roughly 20,000 years ago using a PSMC analysis, followed by a likely explosion of population size starting roughly at the same time as the rise of agriculture in humans. The population size bottleneck distorted the allele frequency distribution, leading to unusually high, but still negative, Tajima’s *D* values, and led to substantially more LD than observed in wild mice. Strikingly, however, we found a very similar pattern in the reduction of *π*/*d* in the tens of kbs flanking protein-coding exons, which are consistent with recurrent selective sweeps on newly arising advantageous mutations.

## Methods

### Samples

We obtained genomic DNA from 22 wild *R. norvegicus* trapped in a ∼500-km^2^ area around the city of Harbin, Heilongjiang Province, China in 2011 (Ness et al., 2012) from locations a minimum of 100 m apart to avoid sampling of closely related individuals. We selected 12 of these individuals for whole-genome sequencing. DNA from one individual black rat (*R. rattus*) that died of natural causes was obtained from Bristol Zoo’s colony.

### Sequencing

Genomic DNA was extracted from a small piece of kidney tissue. Standard Illumina 100bp PE libraries for the HiSeq sequencer with an insert size of approximately 450bp were prepared according to manufacturers recommendations. The Illumina sequencing was performed at the Wellcome Trust Sanger Institute. We obtained a modal coverage of 19x – 46x per sample for *R. norvegicus* and 33x for *R. rattus* (table S4). Reads were aligned to the rn5 reference (from Ensembl release 71) using BWA version 0.5.10-mt Li and Durbin (2009). All lanes from the same library were then merged into a single BAM file using Picard tools (http://picard.sourceforge.net) and PCR duplicates were marked using Picard tools ‘MarkDuplicates’. Finally, the library BAM files were merged into a single BAM containing all sequencing reads for that sample.

### SNP calling and filtering

We used the Genome Analysis Toolkit (GATK) UnifiedGenotyper version 2.8-1-g932cd3a for SNP calling (DePristo et al., 2011), using the following non-default arguments: output mode: emit all confident sites; genotype likelihoods model: both; stand emit conf: 10. By choosing the latter parameter value, we obtained information about sites called with relatively low confidence, which were filtered subsequently, as described below.

Before SNP calling, we first performed indel realignment using GATK IndelRealigner with default parameters on the BAM files containing the aligned reads simultaneously on all 12 samples. There is a SNP database available for *R. norvegicus* from Ensembl, but this contains only 10% of the putative variant sites in our data (estimated from release 71, ftp://ftp.ensembl.org/pub/release-71/variation/vcf/rattus_norvegicus/). Recalibrating bases using a limited SNP data set after realignment may have introduced a significant bias, so we did not do this. It has been shown, moreover, that the combination of local realignment and base recalibration is likely to result in biased SNP calls (Guo et al., 2012). We used all putative indels from a first round of SNP calling to mask all sites near putative indels (for deletions: deleted bases + 1 base on either side; for insertions: insert length + 1 base on either side of insertion point). We also performed a second round of SNP calling using the same parameters (and GATK version 2.7-4-g6f46d11), but without the indel realignment step. We filtered out putative SNPs that did not appear in both sets. Both these steps were done because BWA (and any other alignment algorithm) is prone to introduce false SNPs near indels, particularly in low complexity regions (as described in the online GATK documentation). This occurs because the penalty for introducing a small number of false SNPs may be lower than the penalty for introducing a gap, and is most likely to occur close to sequencing read ends.

We further filtered sites that had a GATK quality score QUAL <23. This threshold was chosen post hoc, based on the distribution of scores of invariant sites. Our samples contained a very small fraction of invariant sites with QUAL <24, above which the density increases markedly (fig. S13). Choosing our threshold just below this value therefore allows filtering on quality without introducing substantial bias against invariant sites. We excluded all sites that had an inbreeding coefficient F<-0.8. GATK only computes F for a site if at least 10 samples are called at that site (online GATK documentation). Using this threshold for F, all sites that have exclusively heterozygous individuals are excluded, which is a strong indication of paralogous reads mapping to the same region. Following common practice, we filtered out high and low coverage regions, since such regions are prone to SNP calling errors. We exploited the fact that we have 12 samples by applying relatively lenient bounds on a per sample basis (between 25 and 300% of the sample’s modal coverage) and using much stricter bounds on the average normalized coverage (between 50 and 140%). The latter bounds were derived from the distribution of autosome wide average coverages (fig. S14). There was more than a factor of two difference between the highest and lowest modal coverage, so we used each individual sample’s modal coverage, computed from the whole autosome, for normalizing coverage. Throughout our analyses, we only considered sites that had at least 3/12 samples called after filtering. In some analyses we applied a “**proximity filter**” that removed all variant sites less than md = 5 bp from another variant site, regardless of site quality. The filter does appear to cause a large number of false negatives, so we applied it only to analyses that are highly sensitive to false positives. The proximity filter had a stronger impact on *π* and D than the precise selection criteria for exons/ CNEs, but had little impact on divergence statistics (figs. S1, S2, S3, S4). It never affected the rank order of different classes of sites.

For the *R. rattus* outgroup we used the same SNP calling pipeline, with the following exceptions: 1) we applied indel realignment to the *R. rattus* genome, aligned to the rn5 reference, in isolation. 2) we used a minimum base quality cutoff of 13 rather than the GATK default 17, based on an analysis of GC content and average base quality using Picard tools CollectGcBiasMetrics. 3) we used a minimum QUAL cutoff of 30, which was determined in the same way as for the *R. norvegicus* data, and is higher because of the higher modal coverage of 33x in the *R. rattus* sample. 4) we masked sites near indels using the *R. rattus* putative indels. 5) we required that sites have a normalized coverage between 40% and 200% of the *R. rattus* modal coverage. We have only one *R. rattus* sample, so filtering against likely paralogs based on Hardy-Weinberg frequencies was not possible.

CpG-prone sites, defined as sites preceded by a C or followed by a G, were identified based on the rn5 and mm10 reference sequences as well at the *R. norvegicus* and *R. rattus* samples. We excluded sites that were CpG-prone in any of these sequences from several of our analyses, because the hypermutability of the CG dinucleotide strongly violates the assumption of a uniform mutation rate across the genome, which is commonly made in the theory underpinning many population genetic analyses. Exclusion of CpG-prone is an effective way of removing this source of hypermutability (Gaffney and Keightley, 2008).

The *R. norvegicus* sample consists of four males and eight females. For consistency in filtering and statistics, we restricted all our analyses to auto-somes, and excluded unplaced contigs.

### Exons

We used the Ensembl Rnor5.0.73 annotation file to obtain the locations of exons. We confined our analysis to exons that are part of complete transcripts, i.e., starting with a start codon, terminated by a stop codon and containing no premature stop codons. The annotation file contains 25,725 transcripts, 20,278 of which are complete: 19,530 on the autosomes, 697 on the X chromosome, 9 on the mitochondrial genome and 42 on unplaced contigs.

Exonic sites were analyzed only if they were consistently 0-fold, 2-fold, or 4-fold degenerate over all annotated transcripts in the rat annotation, based on computational translation of all canonical and non-canonical transcripts in *R. norvergicus* containing the site. Sites with inconsistent degeneracy (e.g., a site that is 4-fold degenerate in a canonical transcript, but 0-fold degenerate in an overlapping non-canonical transcript) were excluded. Confidence intervals were computed using n=1,000 bootstrap replicates by sampling per transcript. For details see supplementary text 1.

### Conserved noncoding elements (CNEs)

Noncoding sequences conserved across the mammalian phylogeny, CNEs, were defined using phastCons on a mammal phylogeny excluding rodents as described in Halligan et al. (2013). This resulted in a set of elements comprising 5% of the genome.

In Halligan et al. (2013), the resulting set of elements was lifted over from human hg18 coordinates to mouse mm9 using liftOver. Each liftOver step inevitably results in the loss of a subset of the elements. Simply lifting over the Halligan et al. (2013) set from mouse mm9 to rat rn5 might, therefore, result in a set biased towards higher conservation. To minimize this potential bias, we used three different routes for lifting over the original hg18 coordinates to our rat rn5 reference: hg18 → hg19 → rn5; hg18 → rn4 → rn5; and hg18 → mm9 → rn5. In case of conflicts, i.e., if different liftOver routes placed the same element at different positions on rn5, we only retained those that differed by no more than 25% in length from the original hg18 element. If different liftOver routes resulted in partially overlapping elements, we retained the one with length closest to the original (fig. S15). From this set of elements, we removed those segments of the elements that overlapped with annotated exons (valid and invalid). This final set of CNEs we call “noOverlap”.

To test how sensitive our results are to the precise definition of CNEs, we also created two slightly different sets of CNEs: “strict”: removing all elements that have any overlap with annotated exons, not just the overlapping segments; and “noOverlap, <1 kb”: the noOverlap set excluding elements longer than 1kb. By analyzing the normalized CNE length distributions for different sets, we found that length distributions are very similar between the human, mouse and rat sets, with a slight inflation of very long elements in the rat (“noOverlap”) and mouse sets (fig. S15). The number of elements in these inflated tails was small, but because the median CNE length is very short, these elements, which are less strongly conserved than the average, could have a disproportionate impact on the within-CNE statistics. From the cumulative length distribution (fig. S15) it can be seen that in the human set, less than 1% of CNE bases occur in elements of over 1kb, whereas in the rat and mouse sets this is 5-6%. For this reason we used the “noOverlap, <1kb” set as default for the within-CNE statistics. For the CNE flanks there was almost no difference between both noOverlap sets, because the number of long (>1 kb) elements is only a very small fraction the total number (fig. S16).

As a neutral reference for the CNEs, we used sequence elements 500 bp upstream and downstream of the CNE, each of half the length of the CNE (Halligan et al., 2013). From this set we masked any segments (noOverlap) or elements (strict) overlapping with exons or other CNEs (from the full noOverlap set).

We used a bootstrapping approach to obtain confidence intervals for the within CNE statistics. We subdivided the genome in 1Mb windows and sampled with replacement among the non-empty windows.

### Estimating the impact of exons on the diversity reduction in CNE flanks

Most CNEs occur in the vicinity of exons. We hypothesized that this explains (part of) the reduction of diversity in CNE flanks, especially farther away from the CNEs. As this effect is likely stronger for CNEs close to exons than CNEs far away from them, we “blurred” the exon flanks based on the distances from CNEs and compared the slopes of the CNE flanks and the “blurred” exon flanks using a linear fit on the data 5-20 kb away from the CNEs.

The idea for the “blurring”, technically called convolution, is taken from image analysis and uses the same principle as simple blurring algorithms in common image manipulation programs. In those, each pixel of image is multiplied with the so called kernel, which determines how far information is smeared out over neighbouring pixels. In our case, we used the 100 bp bins with which the flanks were computed as “pixels”. Instead of a Gaussian kernel, that is typical for image blurring, we used the normalized distribution of distances of CNEs (using the “noOverlap” set) to their nearest exon (fig. S6), up to a maximum of 200 kb, which covers >98.7% of all CNEs.

Before convolution, the exon flanks (not using the proximity filter, i.e., md=0) were padded (extended) with 200 kb of 100 bp bins containing the average value of *π* over the regions 60-100 kb away from the exons in order to accommodate the full width of the kernel.

### Functional annotation

For functional annotation of transcripts, we used the eggNog version 4.0 database, downloaded from http://eggnog.embl.de/version_4.0.beta/downloads.v4.html on 14/01/2014 (Powell et al., 2014).

We combined annotations at different taxonomic levels. In cases of conflicts, we gave preference to annotations of narrower taxonomic levels (e.g. rodents over mammals), with the exception of vague annotations (“R”: “General function prediction only” or “S”: “Function unknown”). We used “X” for transcripts that did not appear in the database. Of the 25,725 transcripts in the annotation file, 21,242 appeared in the database, and of those 16,888 had a function assigned to them (i.e., not R or S). In a subsequent analysis we found that the group of X labeled transcripts was a consistent outlier compared to all other annotations on a variety of metrics. We therefore repeated the within-exon analysis excluding these transcripts to assure that our results do not hinge on this specific subset.

### Genome-wide LD scans

For each variant site (“focal SNP”) we computed the pairwise *r*^2^ statistic directly from the genotypic data (Rogers and Huff, 2009) using up to *n_max_* = 1,500 neighbouring variant sites and sites up to 40 kb away from the focal SNP. We collected averages in bins of 20 bp. The value of *n_max_* was chosen such that it would not lead to the exclusion of sites from the analysis. This value allows for an average diversity of 3.75% within the 40kb, >20 times the average value we found at 4-fold synonymous sites and in 500 bp outside CNEs (fig. 1), before excluding sites. To check this, we verified that the number of sites in the most distant bins did not increase by further increasing *n_max_* in a preliminary analysis using the entire chromosome 19. Note that from the highest diversity estimates reported here, the expected number of SNPs within 40kb would be more than an order of magnitude smaller than our *n_max_*, leaving a wide margin for variation in SNP density. We only considered biallelic sites for which all samples were called and passed the default filters. Moreover, we applied the proximity filter with md=5 for this analysis, because without it we observed small a depression in the 〈*r*^2^〉 curves within 200 bp of the focal SNP, whereas theory predicts a monotonic decline (fig. S8C). This suggests that a fraction of the spurious SNPs that are removed by the proximity filter are of a different kind than the genuine SNPs, which results in a population of sites with much faster decay of 〈*r*^2^〉 than the others. Sites near indels and CpG-prone sites were not considered in this analysis.

### Genome wide LD scans in *M. m. castaneus*

Genotypes were obtained from the VCF files generated by (Halligan et al., 2013). To treat the data as similarly as possible to the rat data, we calculated modal coverage per sample from per sample coverage histograms for the autosome and applied the same bounds as on the *R. norvegicus* data: average normalized coverage between 50 and 140% and per sample between 25 and 300%. We also determined quality histograms for variant and invariant sites, from which we obtained a minimum quality score of 15. Following the original analysis, we disregarded sites with a *χ*^2^ score for HW equilibrium ≥ 0.0002 (against paralogs) and near indels. We removed CpG prone sites based on CpG prone status in *M. m. castaneus*, *M. m. famulus* and *R. norvegicus* (rn4). As with the rat data, all samples had to pass all filters. The mouse sample has a much higher sequence diversity than the rat sample, so to cut computational cost, and because a preliminary analysis showed a much faster decay of 〈*r*^2^〉 than in rat, we only considered sites up to 20 kb away from the focal site. We used *n_max_* = 7,500, equivalent to 15,000 on 40kb or 10x the *n_max_* used for rat, whereas the difference in diversity in regions >80 kb from exons is less than five fold. The proximity filter had virtually no impact on the mouse data (fig. S8C).

### Inference of population history

We used the inference method and software Pairwise Sequentially Markovian Coalescent, PSMC from (Li and Durbin, 2011). For prediction of population size at the most recent time scales, this method is known to be sensitive to false positive heterozygote sites (MacLeod et al., 2013). We therefore required that at least 10/12 individuals were called, so that inbreeding coefficient estimates are available in the GATK output. We applied the proximity filter with md=5 (and md=10 for further sensitivity analysis) and at the individual sample level we only considered calls with a genotype quality ≥ 20. Following (Li and Durbin, 2011), we binned the genome into 100 bp windows and created a fasta-like sequence consisting of the letters “K” (at least one heterozygous site in the bin), “T” (no heterozygous sites) and “N” (less than 10% of sites in the bin called and remaining after filtering). These sequences were then directly input to PSMC using the following arguments: -N80 -r0.63 (for not CpG-prone only) or 1.33 (all sites) -t15 -p “2*4+18*2+4+6”.

These parameters differ from the defaults recommended for the analysis of human data in the PSMC documentation https://github.com/lh3/psmc in three ways. First, we used a much larger number of iterations, after finding that the default (N=25) was insufficient for convergence (convergence of all inferences checked in fig. S17). Second, we reduced the number of free parameters to prevent overfitting (default: -p “4+25*2+4+6”). Third, we initiated r, the ratio *θ*/*ρ*, to per base pair estimates of *μ*/*c* (default: r=5). Parameter t (initial value for history length in units of number of generations/2*N*_0_) was taken from the online documentation, claimed to give good results on human data. The program failed to produce meaningful results using a higher starting value of t=20.

To scale the inferred demographic histories to real time, we assumed a recombination rate of 0.6 cM/Mb (Jensen-Seaman et al., 2004) and a combination of a generation time of 0.5 years (Ness et al., 2012) and a mutation rate of 2.96×10^−9^ per base pair (calculated from our data of divergence from mouse on 4-fold synonymous sites) on the data excluding CpG-prone sites and 5×10^−9^ or 8×10^−9^ on the full sequence data to reflect the higher mutation rate of CpG-prone sites. The latter mutation rate corresponds, for example, to a 2.7 or 5.3 times higher mutation rate, respectively, and CpG-prone sites taking up 40% of the genome and was chosen because it produced either same length genealogies (5×10^−9^) or had the main local minimum coinciding in time (8×10^−9^). As the model assumes that the mutation rate is constant over time and throughout the genome, we considered inferences based on all sequence as less reliable than those based on the non-CpG prone sites, but did produce them as a qualitative control. For details on testing the robustness of the inference, see supplementary text 3.

## Data access

FastQ and BAM files containing all reads aligned to the rn5 reference genome are deposited in the European Nucleotide Archive as study ERP001276 www.ebi.ac.uk/ena/data/view/ERP001276&display=html.

## Acknowledgments

We are grateful to the Wellcome Trust for funding and for grants from the Strategic Priority Research Program of the Chinese Academy of Sciences [XDB11010400 to JXZ], and the China National Science Foundation [91231107 JXZ and 31301887 to YHZ]. We thank David Adams and Thomas Keane for Illumina sequencing and Sue Dow and Adina Valentine (Bristol Zoo) for providing us with a *R. rattus* sample.

